# Effect of a bacterial Glutaminyl Cyclase inhibitor on multi-species-biofilms

**DOI:** 10.1101/2025.10.01.679767

**Authors:** Sigrun Eick, Nadine Taudte, Daniel Ramsbeck, Anna Magdoń, Anton Sculean, Jan Potempa, Mirko Buchholz

## Abstract

Modifying bacterial virulence could be an interesting alternative to antibiotics. The study aimed to examine the effects of an inhibitor targeting bacterial glutaminyl cyclase (which is selectively present in *Porphyromonas gingivalis* (*Pg*), *Tannerella forsythia* (*Tf*), and *Prevotella intermedia* (*Pi*)) on various multispecies biofilms.

Two multi-species biofilms—one containing four species (including *Tf)* and another with 12 species (including *Tf, Pg*, and *Pi*)—were cultured in the presence of 31.25–500 µM of a [4,5-c]pyridine-based inhibitor. After 24 hours, bacterial counts, biofilm mass, metabolic activity, and, when *Pg* was included, Arg-gingipain activity were measured. Additionally, the biofilms were exposed to monocytic cells; here, the release of interleukin (IL)-1β and IL-10 was analyzed. The data were analyzed using a one-way analysis of variance (ANOVA) with a post-hoc comparison performed using the Bonferroni correction.

In all biofilms, total bacterial counts and those of *Pg* and *Tf* remained unaffected by the inhibitor. In the 12-species biofilm, both ‘mass” and total metabolic activity decreased at high inhibitor concentrations (500 µM to 75.2±6.5% and 87.2±5.8%, respectively; each p<0.001). The arginine-specific amidolytic activities of Rgp declined dose-dependently, down to 60.4±10.2% (p<0.001) at 500 µM of the inhibitor. Consequently, *Pg* colonies lost pigmentation as inhibitor concentrations increased. The inhibitor also reduced IL-1β release from monocytic cells stimulated by the 12-species biofilm.

The studied [4,5-c]pyridine-based inhibitor is able to modify virulence of a multispecies biofilm. It might have the potential to be a promising approach in periodontal prevention and therapy.

## Introduction

Periodontitis is a chronic disease that affects the tissues surrounding the teeth and can lead to tooth loss if left untreated. It is one of the most prevalent diseases worldwide; a recent systematic review estimated a prevalence of 61.6% among adult dentate individuals (Trindade et al., 2023). The disease is triggered by a dysbiotic microbiota, which causes an increase in subtypes of epithelial cells and fibroblasts with an inflammatory signature, a dysregulated neutrophil response, an increase in pro-inflammatory (M1) macrophages, and lymphocytes with innate immune properties. All these factors together activate NF-κB signaling, which promotes bone resorption and suppresses the expression of bone matrix proteins (Zhang et al., 2024). The transformation of the eubiotic microbiota into a dysbiotic state associated with periodontal disease has been linked to *Porphyromonas gingivalis*. This keystone pathogen can impair the host’s protective response while promoting inflammation (Hajishengallis and Lamont, 2021). Major virulence factors of *P. gingivalis* include its two arginine-specific (RgpA, RgpB) and one lysine-specific (Kgp) cysteine proteases, also known as gingipains (Hocevar et al., 2018). Other bacterial species, such as *Tannerella forsythia* and *Treponema denticola*, are designated as inflammatory pathobionts, as their virulence genes are upregulated only when tissue destruction has already occurred (Hajishengallis and Lamont, 2021). The subgingival microbiota consists of hundreds of species. Among those enriched in periodontitis compared to periodontal health are, in addition to the previously mentioned three species, *Prevotella intermedia*, various *Fusobacterium nucleatum* subspecies, *Filifactor alocis, Fretibacterium* species, and *Saccharibacteria* TM7 (Abusleme et al., 2021).

According to the European Federation of Periodontology (EFP) S3 level clinical practice guideline (Sanz et al., 2020), the treatment of periodontal disease involves several steps. The first step focuses on controlling supragingival biofilm, gingival inflammation, and risk factors. The second step includes instrumentation, with or without adjuncts. Patients who do not respond adequately receive additional treatment, such as periodontal surgery, in the third step. Finally, all patients should participate in a supportive care program at intervals of 3 to 12 months. As before, controlling supragingival biofilm, gingival inflammation, and risk factors remain essential parts of this stage of therapy (Sanz et al., 2020).

As the dysbiotic biofilm plays a crucial role in disease development, the use of antiseptics and/or antibiotics is discussed in periodontal treatment. According to the guidelines mentioned earlier, the adjunctive use of antiseptics (chlorhexidine) may be considered in the second step of therapy (Sanz et al., 2020). Chlorhexidine is a bisbiguanide and is widely used in dental practice (Thangavelu et al., 2020). It acts quickly to kill many bacteria (Eick et al., 2012). Specifically, when included in a mouthwash, it effectively prevents new biofilm formation (Supranoto et al., 2015). However, adverse effects like tooth discoloration, taste changes, irritation of the oral mucosa or tongue are commonly reported (Supranoto et al., 2015; James et al., 2017).

Using systemic antibiotics as an adjunct improves clinical outcomes (Teughels et al., 2020). However, routine use is not advised; it should only be considered for severe periodontitis in young adults (Sanz et al., 2020). A recent analysis supports this, showing that a combination of amoxicillin and metronidazole effectively halts disease progression, especially in patients with generalized periodontitis at stage III/grade C (Eickholz et al., 2023). Antibiotics can cause side effects like gastrointestinal issues, metallic taste, and nausea (Teughels et al., 2020). Additionally, the rise of antimicrobial resistance has become a growing global concern, with nearly 5 million deaths estimated in 2019 linked to infections caused by antibiotic-resistant bacteria (Collaborators, 2022). One of the key factors contributing to this increase is the widespread use of antibiotics (Chatterjee et al., 2018).

This suggests that antibiotic use should be limited and alternative options explored. Altering bacterial virulence factors, such as gingipains, presents an interesting approach (Pedrosa et al., 2025). One potential target could be glutaminyl cyclase (QC) in *P. gingivalis*. This enzyme is a mammalian-like type II enzyme that catalyzes the cyclization of glutamine (Q) to pyroglutamate (Bochtler et al., 2018; Taudte et al., 2021). Most proteins secreted by *P. gingivalis* via the type 9 secretion system (T9SS), including gingipains, and those exported into the periplasm are transported through the inner membrane via the Sec pathway, where the signal peptide is cleaved by a signal peptidase (Lasica et al., 2017). Downstream of this cleavage site, the proteins often carry a Q, which seems to be cyclized by QC attached to the inner membrane on the periplasmic side (Bochtler et al., 2018). Besides *P. gingivalis*, a QC is also found in *T. forsythia* and *P. intermedia* (Bochtler et al., 2018; Taudte et al., 2021). The first tested QC inhibitor was able to suppress growth in these three species but did not impact other oral bacteria (Taudte et al., 2021). Further development led to the discovery of a new class of *P. gingivalis* QC inhibitors based on a tetrahydroimidazo[4,5-c]pyridine scaffold (Ramsbeck et al., 2021). Selected compounds could inhibit the QC of *P. gingivalis* and *P. intermedia* at concentrations around 0.1 µM, and those of *T. forsythia* at 0.2 µM (Ramsbeck et al., 2021).

The purpose of the experiments described below was to assess the effects of a newly developed [4,5-c]pyridine-based QC inhibitor (S-0636) on two different multispecies biofilms. The main questions were: a) Does the inhibitor affect the overall biofilm structure, including biofilm mass, bacteria, and matrix, as well as total bacterial counts and metabolic activity? b) Does it influence specific bacterial populations, such as commensals and pathobionts like *T. forsythia, P. gingivalis, P. intermedia*? c) Does it impact bacterial virulence, particularly the Arg-gingipain activity of *P. gingivalis*? d) What concentration of the inhibitor is needed to disrupt the biofilm? e) Are there differences between the biofilms composed of 4 species without *P. gingivalis* and those with 12 species? and f) Does pretreating the biofilm with the inhibitor alter the immune response? The purpose of the experiments outlined below was to determine the effects of a newly developed [4,5-c]pyridine-based QC inhibitor (S-0636) on two different multispecies biofilms. The key questions were: a) Does the inhibitor impact the biofilm overall structure, including biofilm mass covering bacteria and matrix, total bacterial counts, and metabolic activity? b) Does it influence specific bacterial populations (such as commensals and pathobionts like *T. forsythia, P. gingivalis, P. intermedia*)? c) Does it affect bacterial virulence, specifically the Arg-gingipain activity of *P. gingivalis*? d) What concentration of the inhibitor disrupts the biofilm? e) Are there differences between the different types of biofilms (4-species without *P. gingivalis*, and 12-species)? and f) Does pretreating the biofilm with the inhibitor modulate the immune response?

## Materials and methods

### Inhibitors

From a panel of different compounds, a [4,5-c]pyridine-based inhibitor was selected: S-0636. Its synthesis is described elsewhere (Taudte 2025, submitted). It was dissolved in phosphate buffered saline (PBS, Gibco™ PBS pH 7.4, Thermo Fisher Scientific, Waltham, MA, USA). The inhibitor was used at final concentrations of 500 µM, 250 µM, 125 µM, 62.5 µM, and 31.25 µM in the assays. PBS served as a negative control.

### Microorganisms in a defined biofilm

Two different biofilms were used. The four-species biofilm included *Streptococcus gordonii ATCC 10558, Actinomyces naeslundii* ATCC 12104, *Fusobacterium nucleatum* ATCC 25586, and *Tannerella forsythia* ATCC 43037. For the 12-species biofilm, the following species were additionally included: *Porphyromonas gingivalis* ATCC 33277, *Prevotella intermedia* ATCC 25611, *Campylobacter rectus* ATCC 33238, *Capnocytophaga gingivalis* ATCC 33624, *Eikenella corrodens* ATCC 23834, *Filifactor alocis* ATCC 33238, *Parvimonas micra* ATCC 33270, and *Treponema denticola* ATCC 35405.

The strains were passaged on tryptic soy agar plates (Oxoid, Basingstoke, GB) with 5% sheep blood (and with 10 mg/l N-acetylmuramic acid for T. forsythia). T. denticola ATCC 35405 was maintained in modified mycoplasma broth (BD, Franklin Lakes, NJ) enriched with 1 g/ml cysteine and 5 μg/ml cocarboxylase. All strains were cultured at 37°C: streptococci and A. naeslundii ATCC 12104 in 10% CO_2_, and the other strains under anaerobic conditions.

### Biofilm formation

Ninety-six-well plates were coated with 10 µl per well of a protein solution containing 0.67% mucin (Merck KGaA, Darmstadt, Germany) and 1.5% bovine serum albumin (SERVA Electrophoresis GmbH, Heidelberg, Germany) for 15 minutes. Bacterial strains were suspended in a 0.9% w/v NaCl solution based on OD600nm=1. Then, the bacterial mixture was prepared using 1 part *S. gordonii*, 2 parts *A. naeslundii*, and 4 parts of the other strains except *T. forsythia, P. gingivalis*, and *T. denticola*. This mixture was added at a ratio of 1:9 to the culture medium (Wilkins-Chalgren broth (Oxoid, Basingstoke, GB) with 10 mg/l β-NAD and 10 mg/l N-acetylmuramic acid). From this suspension, 200 µl was pipetted into each well, and the plate was incubated for 6 hours under anaerobic conditions. Afterward, 50 µl of a suspension (prepared as above for the other strains) containing *T. forsythia* (*P. gingivalis* and *T. denticola*, according to the experiments), along with the inhibitor at the respective concentration or a control, was added. Following an additional 18 h of incubation (total 24 h) in anaerobic conditions, the samples were analyzed.

### Analysis of biofilms

The supernatant of the biofilm was removed, and the biofilms were briefly and carefully washed. Then, 250 µl of 0.9% w/v NaCl was added. The biofilms were scraped from the surface and thoroughly mixed. From that suspension, 25 µl were used to determine colony-forming units (CFU), 25 µl for measuring metabolic activity, 25 µl for biofilm mass determination, 25 µl for qPCR, and 100 µl for quantifying the arginine-specific amidolytic activity (BApNA).

A 10-fold dilution series was performed on the aliquot designated for CFU determination. Each 25 µl sample was plated on tryptic soy agar plates (Oxoid) supplemented with 5% sheep blood. One series was incubated under anaerobic conditions, representing total CFU counts, while the other was incubated with 10% CO_2_ to count streptococci and actinomyces (as commensals). For measuring metabolic activity, 200 mg/l resazurine (Merck KGaA) was mixed with nutrient media at a ratio of 1:10, and 100 µl per well was added to the biofilm suspension in a 96-well plate. After a 1-hour incubation at 37°C, the plate was read at 570 nm against 600 nm. For quantifying the biofilm mass, biofilm aliquots were pipetted into a new 96-well plate and fixed on the surface at 60°C for 60 minutes. Next, 50 µl of a 0.06% (w/v) crystal violet solution was added per well. After washing, 50 µl of 30% acetic acid was added, and after 10 minutes, the plate was read at 600 nm.

The counts of *T. forsythia, P. gingivalis*, and *P. intermedia* were determined by using qPCR (Jentsch et al., 2020). Additionally, in a 12-species biofilm (including *P. gingivalis*), the arginine-specific amidolytic activity was measured. The respective suspension was centrifuged for 10 min with 5000 *g* at 20°C. To the sediment, 200 µl of NaCl was added, and the mixture was exposed to ultrasonication, ensuring that cell-wall activity was also included in the measurements. The amidolytic activity was monitored at 405 nm for 1 h at 37°C after adding 0.5 mM N-a-benzoyl-DL-arginine-p-nitroanilide (BApNA; Merck KgaA) in 1.0 ml of 0.2 M Tris– HCl, 0.1 M NaCl, 5 mM CaCl2, 10 mM cysteine, pH 7.6.

Additionally, in selected samples (12-species biofilm, 500 µM of inhibitor and control), the expression of gingipains was measured, and the location of P. gingivalis and T. forsythia was visualized. To quantify gingipain expression, RNA was extracted, cDNA was synthesized, and qPCR was performed as described previously (Eick et al., 2019). For visualization by FISH technology, biofilms were cultured as before but on glass slides in 24-well plates. Samples were fixed in 4% paraformaldehyde / PBS for 1 h. After washing in PBS and preincubating in hybridization buffer (20% formamide, 0.9M NaCl, 20 mM Tris-HCl, 0.01% SDS) at 46°C for 15 min, the probes for P. gingivalis (Pg-Cy3 (Sunde et al., 2003)), T. forsythia (Tf-Atto-488, sequence (Sunde et al., 2003)), and all bacteria (EUB 338-Atto-425, sequence (Amann et al., 1990)) at concentrations of 1 µM, 1.5 µM, and 0.5 µM were added to the hybridization buffer and incubated for 3 h at 46°C. After several washing steps, samples were embedded and visualized using a Zeiss LSM 710 confocal microscope (Carl Zeiss). Additional visualization was conducted with Imarisviewer software (Bitplane, IMARIS 10.0.0).

### Interaction of monocytic cells with biofilm

The biofilms were formed as described above for 24 h.

MONO-MAC-6 cells (DSMZ no. ACC 124), a monocytic cell line of human origin, were cultured with RPMI 1640 medium (Invitrogen; Carlsbad, CA, USA) supplemented with 10% fetal bovine serum (FBS; Invitrogen). Before use in experiments, MONO-MAC-6 cells were washed once and adjusted to 10^^5^ cells/mL in RPMI 1640 medium with 0.5% FBS and the inhibitor at the respective concentration. Please note there is always a cell control without biofilm.

The supernatants from the biofilms were carefully removed and replaced with 200 µl per well of MONO-MAC-6 cell suspension, with or without inhibitor. After 4 h of incubation, the media were taken out, centrifuged at 250 *g* at 20°C for 10 min. The supernatants were stored at −80°C until analysis. Commercial ELISA kits (R&D Systems, Minnesota, MN, USA) were used to measure the levels of released interleukin IL-1β and IL-10 according to the manufacturer’s instructions. The detection limit was 1 pg/ml for both cytokines.

An influence on cell vitality by the experimental settings was screened using the MTT assay according to Mosmann (Mosmann, 1983).

### Statistical analysis

In all biofilm experiments, each of eight independent biological samples (obtained in two series) was included in the statistical analysis. Bacterial counts were recorded as log10. In presentations, the arbitrary units of activity and the mass data of the biofilm were related to the means of the untreated controls. The data were compared using a one-way analysis of variance (ANOVA) with a post-hoc comparison employing Bonferroni correction. The statistical analysis was performed with SPSS 29.0 (IBM, Armonk, NY, USA).

## Results

Of the post-hoc analyses, only statistical differences vs. control are presented.

### Four-species biofilm

The total CFU counts of the 4-species biofilm without the addition of an inhibitor were 7.42±0.10 log10, and the counts for T. forsythia were 6.65±0.15 log10. There was no statistically significant difference in bacterial counts, biofilm activity, or biofilm mass when any concentration of the inhibitor was added (Fig. 1).

**Fig. 1:**
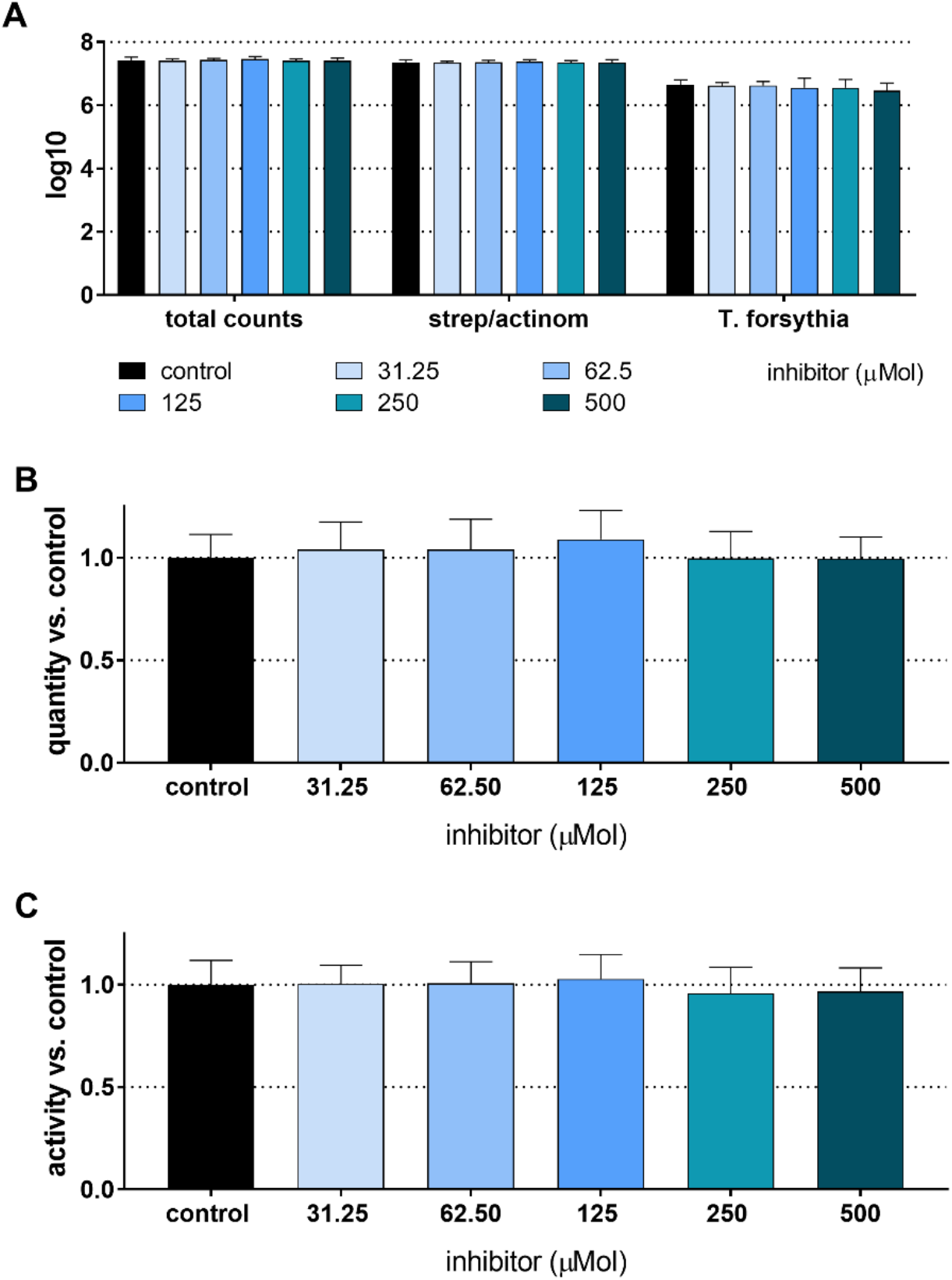
Total counts and counts of streptococci/actinomyces and Tannerella forsythia (A), mass (B), and metabolic activity (C) of 24 h 4-species biofilms cultured with and without different concentrations of S-0636.

**Fig. 2:**
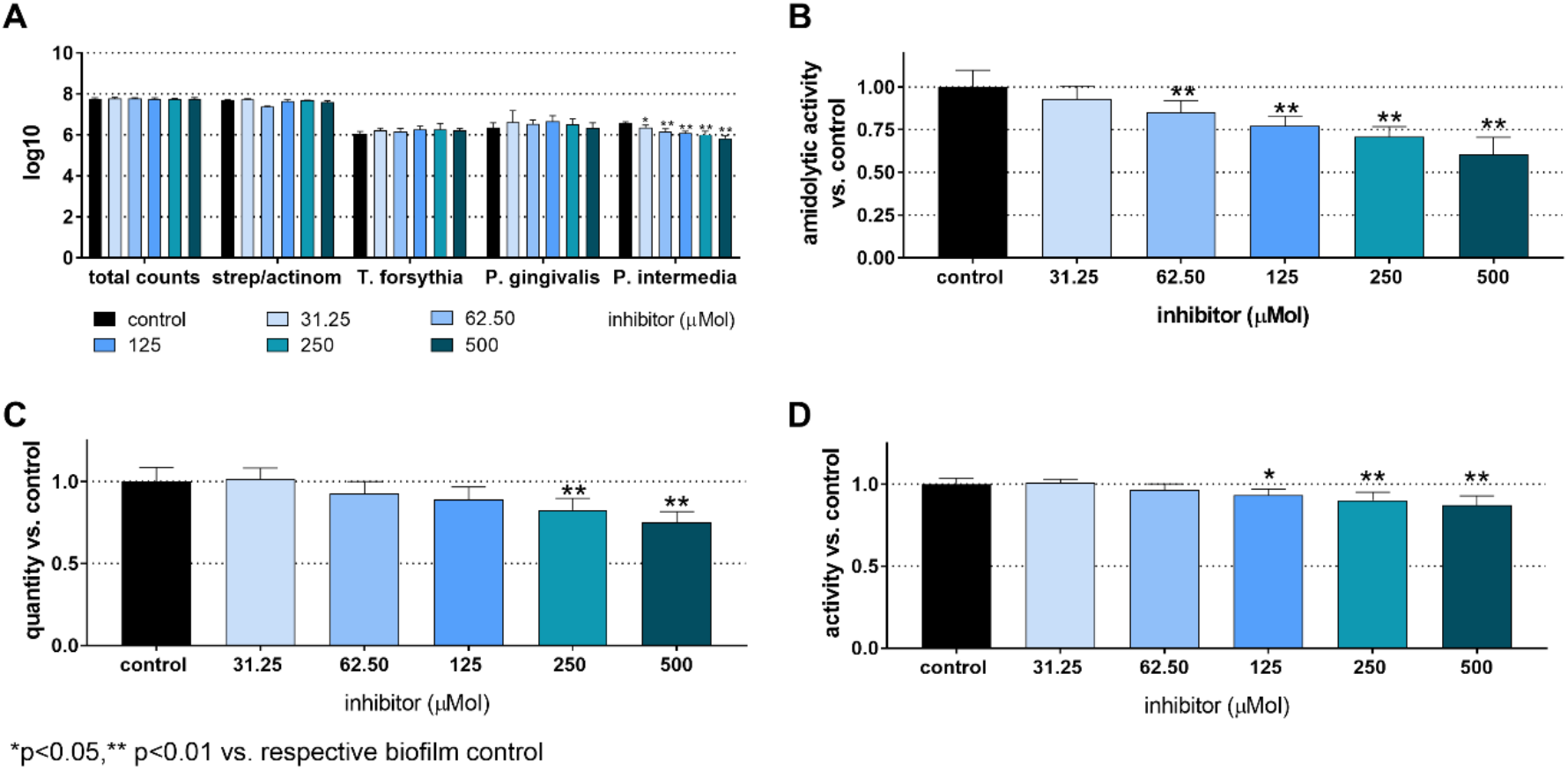
Total counts and counts of streptococci/actinomyces, *Tannerella forsythia, Porphyromonas gingivalis*, and *Prevotella intermedia* (A), arginine-specific amidolytic activity (B), mass (C), and metabolic activity (D) of 24 h 12-species biofilms cultured without and with different concentrations of S-0636. ^*^ p<0.05, ^**^p<0.01 vs. control

### Twelve-species biofilm

The total CFU counts in the 12-species biofilm without inhibitor added were 7.75±0.05 log10. *T. forsythia* counts were 6.05±0.12 log10, P. gingivalis counts were 6.37±0.23 log10, and *P. intermedia* counts were 6.55±0.09 log10. No bacterial counts showed a statistically significant difference, except for *P. intermedia*, which decreased in a concentration-dependent manner. The lowest counts occurred at 500 µM of the inhibitor (5.84±0.11 log10). The biofilm “mass” was significantly reduced when treated with 250 µM (82.7±7%, p<0.001) and 500 µM (75.2±6.5%, p<0.001) of the inhibitor. Additionally, the total biofilm metabolic activity decreased significantly at inhibitor concentrations of 125 µM and higher (125 µM: 93.3±3.7%, p=0.033; 250 µM: 90.3±4.8%, p<0.001; 500 µM: 87.2±5.8%, p<0.001). The arginine-specific amidolytic activities of the biofilm decreased in a concentration-dependent manner with the inhibitor, and the results were statistically significant compared to the control at 62.5 µM (85.2±7.0%, p=0.009), 125 µM (77.1±5.9%, p<0.001), 250 µM (70.7±6.2%, p<0.001), and 500 µM (60.4±10.2%, p<0.001).

It is interesting to note that *P. gingivalis* colonies lost pigmentation after exposure to high concentrations of the inhibitors (Fig. 3).

**Fig. 3:**
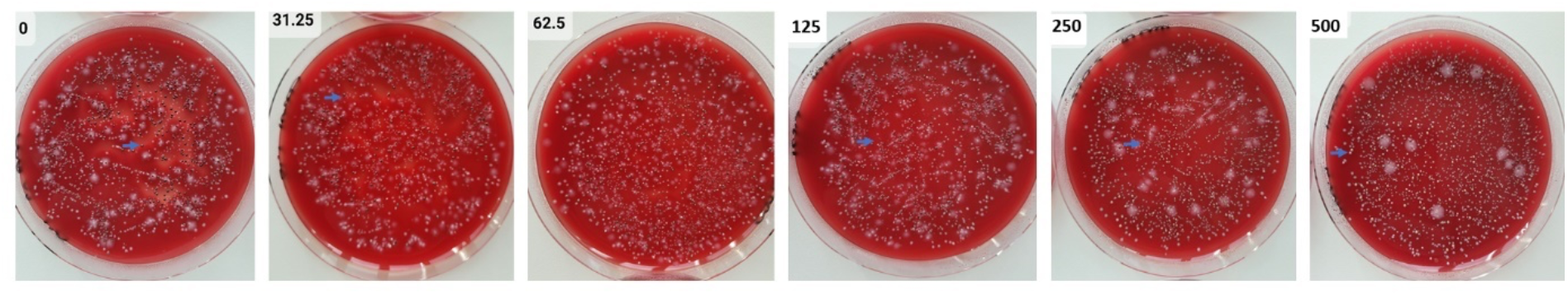
Agar plates with samples from 24 h 12-species biofilms were tested both without and with varying concentrations of S-0636. After performing a dilution series, aliquots of the biofilms were spread on agar plates and incubated anaerobically for 8 days. *P. gingivalis* (→) forms black colonies without the inhibitor; with increasing concentrations of the inhibitor, its colonies gradually lose their pigmentation.

### mRNA expression of gingipains and FISH staining of biofilms

The analysis of mRNA expression of the gingipains after culturing the biofilm with 500 µM of the inhibitor showed almost no change for kgp (0.84±0.17-fold) and a slight increase for rgpA (1.81±0.47-fold) and rgpB (3.54±0.17-fold) compared to the untreated biofilm. Confocal laser scanning microscopy (CLSM) images were taken from biofilms stained with three different FISH probes. These images confirm the greater thickness of the control biofilms compared to the biofilms exposed to 500 µM of the [4,5-c]pyridine-based inhibitor (Fig. 4).

**Fig. 4:**
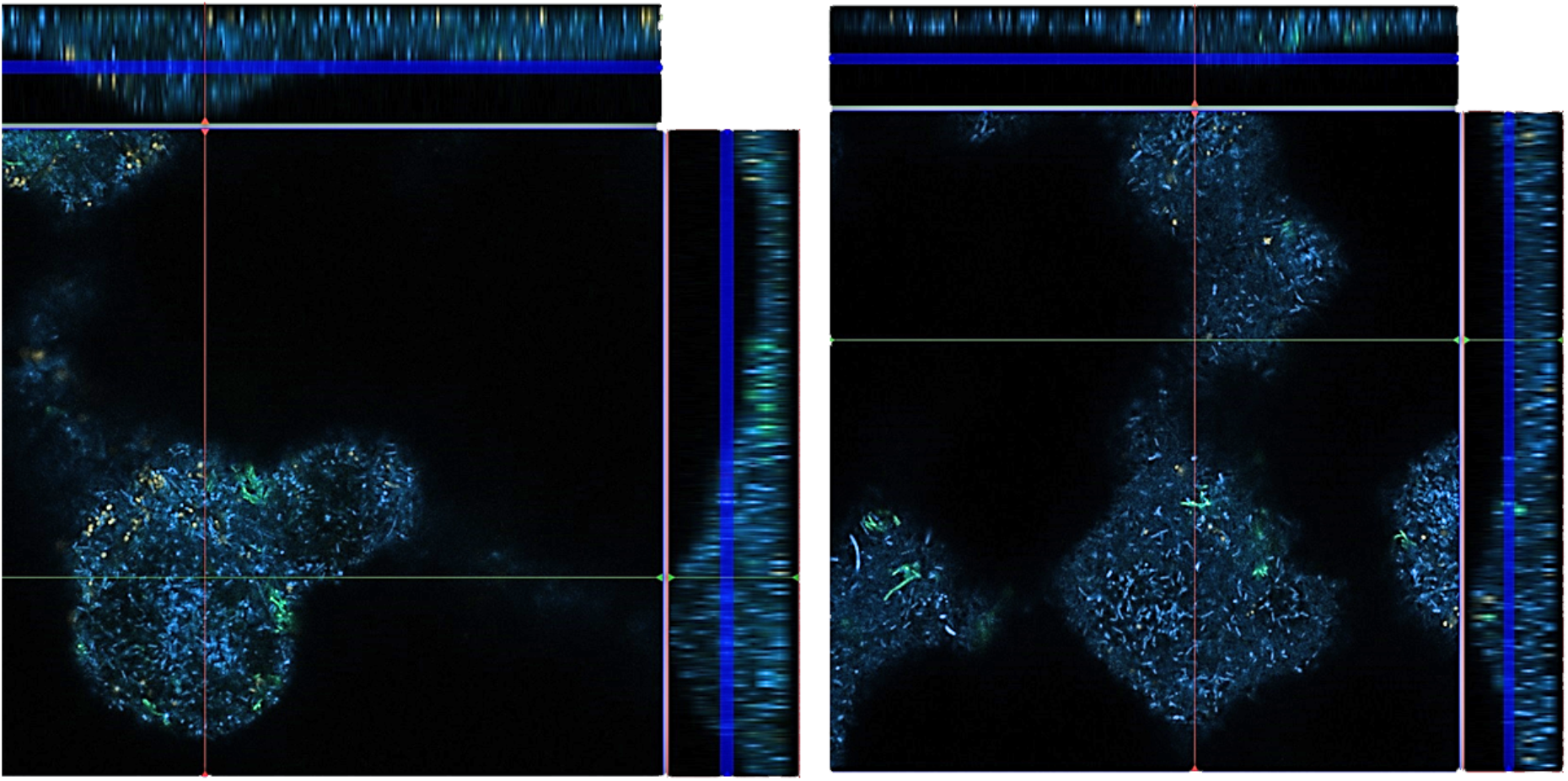
FISH images of 12-species biofilms cultured without (left) and with 500 µM of S-0636 (right). *P. gingivalis* appears yellow-orange, *T. forsythia* green, and all other bacteria are blue.

The photographs taken from different layers of the biofilm suggest that *P. gingivalis* is primarily located at the bottom, *T. forsythia* is found more in the middle layers of the biofilm, and it appears that *T. forsythia* forms clusters within the biofilms. Clear differences between the control biofilm and those cultured with 500 µM of the inhibitor are not visible (Fig. 5).

**Fig. 5:**
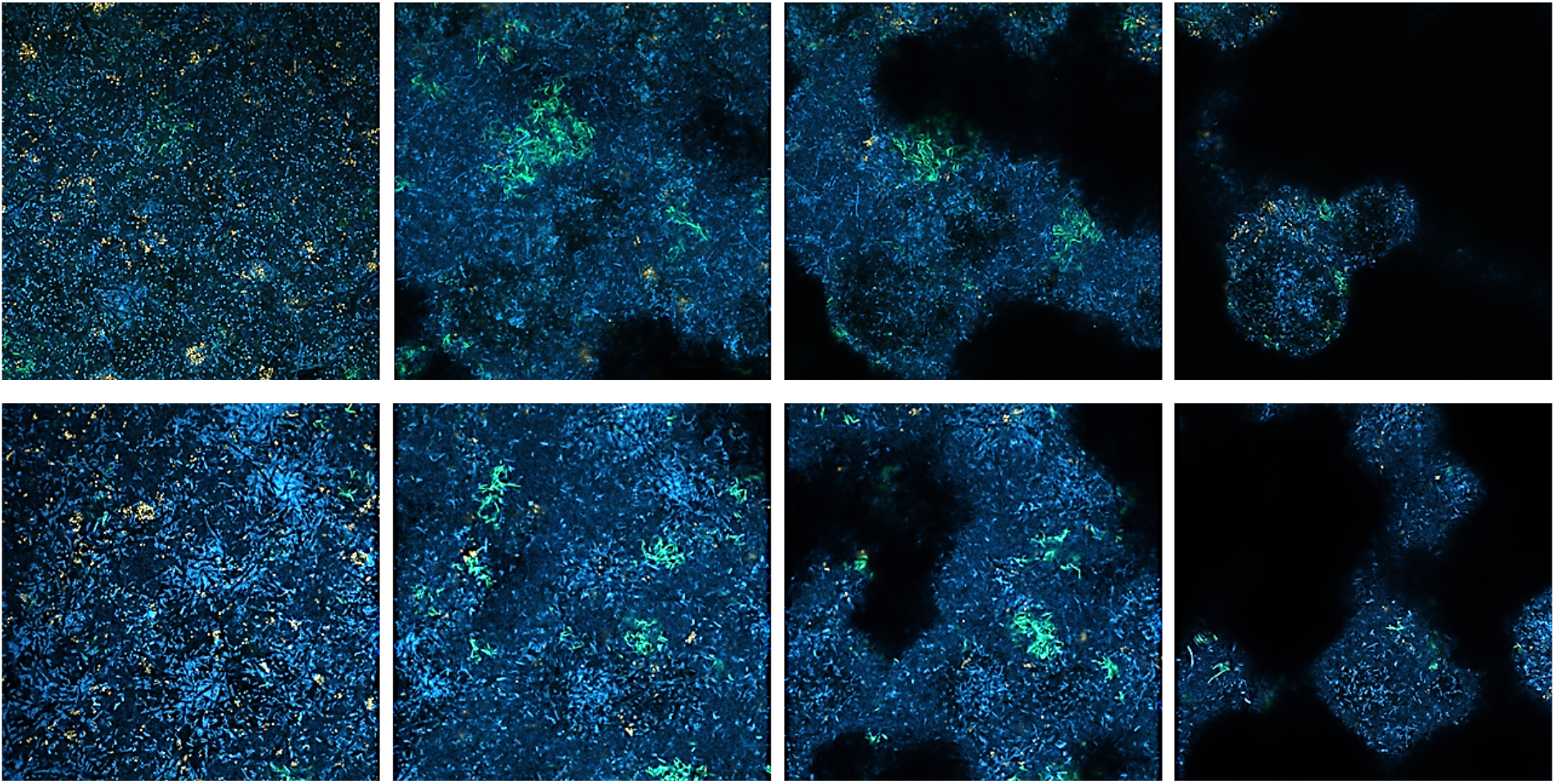
FISH images of 12-species biofilms cultured without (upper row) and with 500 µM of the [4,5-c]pyridine-based inhibitor (lower row). Different layers of the biofilm from bottom to top (left to right) are visualized. P. gingivalis appears yellow-orange, T. forsythia green, and all other bacteria are blue.

### Release of IL-1β and IL-10

Without bacterial stimulus, the MONO-MAC-6 cells did not release detectable IL-1β, but they did produce moderate amounts of IL-10. With the influence of biofilms, IL-1β levels increased and were highest in the case of the 12-species biofilm without inhibitor. The [4,5-c]pyridine-based inhibitor reduced IL-1β release stimulated by the 12-species biofilm. In the case of the 4-species biofilm, IL-1β levels in the presence of the compound did not change significantly compared to the 4-species biofilm without inhibitor (Fig. 6A). Exposure to biofilms (both 4- and 12-species) decreased IL-10 levels. The inhibitor had no additional effect (Fig. 6B). The MTT test confirmed that S-0636 did not cause any cytotoxic reaction in the MONO-MAC-6 cells.

**Fig. 6:**
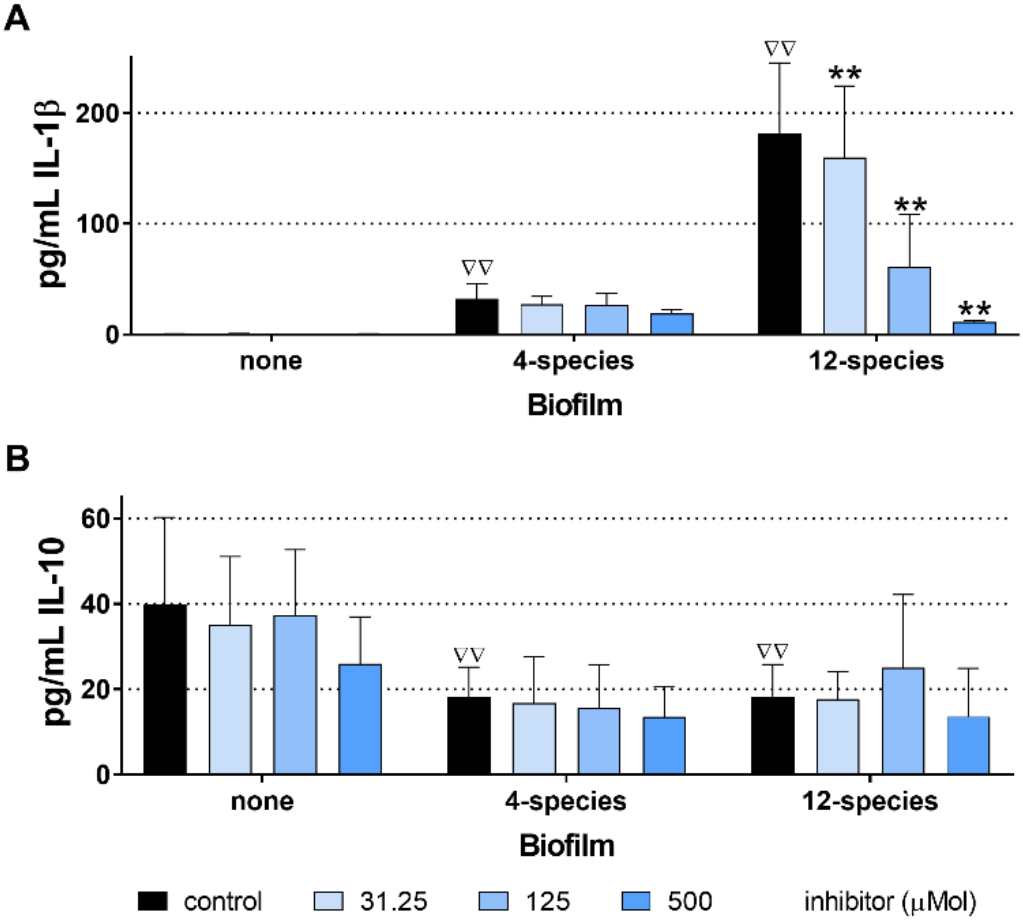
Levels of interleukin (IL)-1β (A) and IL-10 (B) released from MONO-MAC-6 cells after 4 h exposure to biofilms cultured without and with various concentrations of a [4,5-c]pyridine-based inhibitor, ΔΔp < 0.01 compared to no biofilm (comparison only made for controls).^**^p<0.01 vs. respective biofilm control

## Discussion

The experiments focused on the effect of a [4,5-c]pyridine-based QC inhibitor on two defined biofilms. Two biofilms were included: a 4-species biofilm containing bacteria with a QC-only *T. forsythia*, and a 12-species biofilm that also included *P. gingivalis* and *P. intermedia*.

One of the questions to be answered was whether there was a difference between the two biofilms. The 4-species biofilm was not affected by the inhibitor. Statistically significant differences were found only when using the 12-species biofilm. It might be linked to a stronger inhibition (lower ki-value) of *P. gingivalis* QC compared to *T. forsythia* QC (Jäger et al. 2019).

Other questions were whether the inhibitor affects the biofilm as a whole and whether it alters the counts of specific bacteria. The total bacterial counts in the biofilm did not change. The counts of individual species, including *P. gingivalis, T. forsythia*, and the commensals (streptococci/actinomyces), remained stable. This is supported by the FISH images, which clearly showed no differences in the counts of *P. gingivalis* and *T. forsythia*.

Only *P. intermedia* showed a dose-dependent decrease in the number of cells in the biofilm treated with the inhibitor. A direct bactericidal effect of the inhibitor via QC can likely be ruled out, as the inhibitor’s potency was higher compared to *P. gingivalis* QC (Jäger et al., 2019). One possible reason for the reduction in *P. intermedia* numbers in the biofilm may be a decrease in the coaggregation ability between *P. gingivalis* and *P. intermedia*. An important coaggregation factor of *P. gingivalis* with *P. intermedia* is a protein from the outer membrane vesicles encoded by gingipain genes (Kamaguchi et al., 2003), whose secretion can be hindered.

The lower metabolic activity and biofilm mass could indicate impaired synergy in biofilm formation. Since total bacterial counts were unaffected, the reduced biofilm mass suggests an effect on the biofilm’s matrix. Synergistic effects among periodontal bacteria have been described for *P. gingivalis*. Diffusible signaling molecules from *P. gingivalis* can interfere with *F. nucleatum*’s metabolism and promote increased polysaccharide synthesis (Yamaguchi-Kuroda et al., 2023). A strong synergy in biofilm formation was observed for *P. gingivalis* and *T. denticola*, with gingipains playing a key role (Zhu et al., 2013). RgpA appears to facilitate the growth of other oral species within a community, including *P. gingivalis, P. micra, F. nucleatum*, and *Streptococcus constellatus* (Davies et al., 2021). Considering the specificity of the inhibitor, molecules transported by the T9SS may also contribute to the formation of the biofilm matrix. According to functional categories, a high percentage of proteins involved in T9SS, outer membrane functions, and peptidoglycan biosynthesis carry an N-terminal Q (Szczesniak et al., 2023). Pyroglutaminylation of these proteins by QC appears to stabilize them and protect them from degradation (Szczesniak et al., 2023). It can be assumed that fewer proteins contributing to matrix formation are available; however, knowledge about proteins produced by *P. gingivalis, T. forsythia*, and *P. intermedia* that contribute to biofilm matrix synthesis remains limited.

The Arg-gingipain specific amidolytic activity, which correlates with *P. gingivalis* virulence, was clearly reduced by the QC inhibitor in a concentration-dependent manner. Our data suggest limited synthesis or processing of gingipains. However, limited synthesis can be ruled out at least at the mRNA level. Synthesized prepro-gingipains undergo extensive post-translational proteolytic processing, similar to many other virulence factors, as they are first transported by the Sec system to the periplasm and then through the T9SS (Veith et al., 2022). Blocking specific components of T9SS caused a loss of cell-associated gingipain activity (Kgp or total) and colony pigmentation (Heath et al., 2016; Naito et al., 2019). In our study, we also observed a loss of colony pigmentation in *P. gingivalis*. It has been shown that glutamine cyclization into pyro-glutamate plays a role in the cell-associated activity of RgpA (Bochtler et al., 2018). The underlying mechanisms might be explained by the fact that proteins with N-terminal Q involved in T9SS (Szczesniak et al., 2023) are either not functional or only partially functional when a QC inhibitor is present.

Furthermore, we evaluated the potential interaction between the QC inhibitor’s effect on the biofilm and the host response. For this, the pre-cultivated biofilm, with and without the inhibitor, was exposed to monocytic cells for 4 hours, after which the release of interleukin (IL)-1β, a pro-inflammatory cytokine, and IL-10, an anti-inflammatory cytokine, was measured. In periodontitis, macrophages can be polarized into M1 or M2 types. M1 macrophages, which promote inflammation, are characterized by high levels of pro-inflammatory cytokines, such as IL-1. In contrast, M2 macrophages, which facilitate anti-inflammatory functions, are marked by high levels of IL-10 (Sun et al., 2021). The initiation and progression of periodontal disease are driven by a dysregulated inflammatory response to a dysbiotic biofilm (Nguyen et al., 2020). This is supported by *in vitro* results showing high levels of IL-1β and sustained levels of IL-10 after exposure of the 12-species biofilm to monocytic cells. Clinically, high IL-1β levels decrease after periodontal therapy, coinciding with improvements in periodontal clinical parameters (Nanakaly et al., 2024). In the present study, IL-1β levels also decreased following the application of the inhibitor.

Results in the 12-species biofilm were clearly dependent on the concentration of the inhibitor used. Since the compound will be incorporated into an oral health-care product for local application, it should be available at a sufficiently high concentration. It should be noted that in this in vitro study, the concentration of the inhibitor remained stable within the experimental setup. In vivo, depending on the application site, the flow of saliva or gingival crevicular fluid must be considered. The mean unstimulated salivary flow rate was measured at 287 µl/min (Gill et al., 2016), while stimulated salivary flow can reach up to 640 µl/min (Ligtenberg et al., 2020). Gingival crevicular fluid flow was approximately 0.5 µl/min (Lima et al., 2018). To address the issue of immediate dilution of the compound, it may be necessary to incorporate it into a slow-release device that can maintain the active concentration over an extended period (Schmid et al., 2020).

The advantage of the study was using a clearly defined study design that allowed comparison of the two biofilms and different inhibitor concentrations. The studied variables enabled conclusions about the anti-biofilm, anti-virulence, and immunomodulatory properties of the inhibitor. However, the models only reflected part of the situation in the oral cavity; the biofilm consisted of up to 12 species, the interaction with host cells included only a monocytic cell line, and salivary or gingival crevicular fluid flow was not considered. These could be limitations of the study, highlighting the need for further in vitro research with more complex models. Finally, the efficacy of the inhibitor must be demonstrated through randomized clinical trials.

Overall, the compound has potential for inclusion in oral health care products. It may alter the virulence of a dysbiotic biofilm, leading to a balanced pro- and anti-inflammatory response. Since the inhibitor clearly acts in a concentration-dependent manner, it should be included at a sufficiently high concentration in an oral health care product.

## Data availability statement

The raw data supporting the conclusions of this article will be made available by the authors, without undue reservation.

## Authors contributions

SE: Conceptualization, Methodology, Data curation, Supervision, Writing - original draft, review and editing, NT: Conceptualization, Writing - original draft, review and editing,, DR: Conceptualization, Writing - review and editing AM: Investigation, Formal analysis, Writing – review and editing, AS: Resources, Writing - review and editing, JP: Conceptualization, Writing - review and editing, MB: Funding acquisition, Conceptualization, Writing - review and editing

## Generative AI statement

The authors declare that no Gen AI was used in the creation of this manuscript.

## Acknowledgement

We are grateful to Xilei Zhu for help in taking the CLSM photographs.

